# Estimating the genome-wide mutation rate from thousands of unrelated individuals

**DOI:** 10.1101/2022.07.11.499645

**Authors:** Xiaowen Tian, Ruoyi Cai, Sharon R. Browning

**Author notes:** Corresponding authors: X. Tian, S. R. Browning.

## Abstract

We provide a method for estimating the genome-wide mutation rate from sequence data on unrelated individuals by using segments of identity by descent (IBD). The length of an IBD segment indicates the time to shared ancestor of the segment, and mutations that have occurred since the shared ancestor result in discordances between the two IBD haplotypes. Previous methods for IBD-based estimation of mutation rate have required the use of family data in order to accurately phase the genotypes. This has limited the scope of application of IBD-based mutation rate estimation. Here, we develop an IBD-based method for mutation rate estimation from population data, and we apply it to whole genome sequence data on 4,166 European American individuals from the TOPMed Framingham Heart Study, 2,966 European American individuals from the TOPMed My Life Our Future study, and 1,586 African American individuals from the TOPMed Hypertension Genetic Epidemiology Network study. Although mutation rates may differ between populations due to genetic factors, demographic factors such as average parental age, and environmental exposures, our results are consistent with equal genome-wide average mutation rates across these three populations. Our overall estimate of the average genome-wide mutation rate per 10^8^ base pairs per generation for single nucleotide variants is 1.24 (95% CI 1.18-1.33).

## Introduction

Although the genome-wide average human mutation rate is a critical parameter in genetic studies, there is still considerable uncertainty about its value. Sequence data on parent-offspring trios is increasingly available and used to estimate the mutation rate,^1–3^ however genotype error rates are high relative to rates of mutation. Thus, the choice of filtering criteria to balance false positive and false negative genotype calls is critical and can significantly impact the estimates.^4^ Sequence data on multi-generational families can help address these issues,^2^ but are unavailable for many populations.

Furthermore, differences in mutation rates between human populations are likely, although they may not be large. Mutation rates increase with parental age,^5; 6^ and average parental age can vary between populations. Differing environmental conditions and exposures to mutagens, as well as genetic factors, may also lead to differences between populations.^3^ Thus, estimates of average genome-wide mutation rate by population are needed, rather than simply obtaining a single “human” rate.

Recently, methods have been developed for estimating mutation rates from segments of identity by descent (IBD).^7–10^ Existing methods require highly accurate estimates of haplotype phase,^8; 10^ except for methods that utilize homozygosity by descent (i.e. IBD within an individual rather than between individuals).^7; 9^ In particular, highly accurate phase is needed for the rarest variants, because these are the variants that are the most likely to be due to mutation since the common ancestor of the IBD haplotypes. However, statistical population-based phasing is unable to provide accurate phase of variants that are seen only a few times in the sample.^11^ Thus, previous IBD-based methods used data from parent-offspring trios, for which Mendelian rules enable phasing of rare variants.^8; 10^

In this paper we build on the work of Tian et al.,^10^ which used sets of three IBD haplotypes to estimate the mutation rate. By focusing on sets of three IBD haplotypes, the impact of genotype error is much reduced. Variants must be seen at least twice to be considered, and must be carried by two of the three IBD haplotypes, greatly increasing their likelihood of being true rare alleles rather than mis-calls. We show that this design also enables highly accurate phasing of the rare variants via IBD, so that parent-offspring trios are not needed. As a result, our method can be applied to sequenced population samples, such as those in the TOPMed study.^12^

Existing IBD-based methods adjust for gene conversion with post-hoc regression,^8–10^ but in this work we incorporate gene conversion directly within the likelihood framework.

## Methods

### Modelling of mutations, gene conversion, and genotype error

We model the occurrence of mutations as a Poisson process of rate *μ* per base pair per meiosis. Thus, across *g* independent meioses and *l* base pairs the number of changed alleles due to mutation is distributed as Poisson with mean *lgμ*.

The lowest frequency variants are useful for estimating the mutation rate, while the highest frequency variants are most informative for estimating IBD segments. We thus divide the genetic markers into three groups: markers with minor allele frequency (MAF) < *f*_1_, which are informative about mutation and gene conversion; markers with MAF between *f*_1_ and *f*_2_, which can be affected by gene conversion but not by mutation, thus allowing estimation of gene conversion rates for correcting for gene conversion in the markers with MAF < *f*_1_; markers with MAF > *f*_2_, which are used for IBD segment detection. Gene conversions can disrupt identity by state, leading to inability to detect the corresponding IBD segments. By using distinct sets of markers for IBD detection and parameter estimation we avoid this cause of downward bias in estimates of gene conversion rates. We use *f*_1_ = 0.1 and *f*_2_ = 0.25 in all analyses. In simulation studies, we found that all mutations in IBD segments had allele frequencies < 0.1, and using *f*_2_ = 0.25 allows for both sufficient markers for estimation of gene conversion rates and sufficient markers for IBD detection.

Let *θ* be the proportion of base pairs that are located within a gene conversion tract per meiosis. The offspring’s haplotype is only altered by gene conversion at positions where the parent individual’s genotype was heterozygous. We write *h*(0, *f*_1_) for the genome-wide proportion of base pair positions that are heterozygous and have MAF < *f*_1_ in the population. We estimate *h*(0, *f*_1_) from the genome-wide data. For a single base pair position and meiosis, the expected number of alleles with frequency < *f*_1_ inserted by gene conversion will be *θh*(0, *f*_1_)/2. The division by 2 is because if the individual in which the gene conversion occurred was heterozygous at a locus within the gene conversion tract, either the transmitted haplotype was originally the major allele and changed to the minor allele by the gene conversion, or vice versa, each with probability 1/2. We are only counting instances of change to the minor allele here. Thus, we model the number of alleles with frequency < *f*_1_ inserted by gene conversion over *g* meioses and *l* base pairs as having a Poisson distribution with mean *lgθh*(*f*_1_, *f*_1_)/2. Similarly, we model the number of alleles with frequency >1 – *f*_1_ inserted by gene conversion over *g* meioses and *l* base pairs as having a Poisson distribution with mean *lgθh*(0, *f*_1_)/2, using the same reasoning but this time only counting instances in which the transmitted haplotype’s allele was changed from the minor to the major allele.

We write *h*(*f*_1_, *f*_2_) for the genome-wide proportion of base pair positions that are heterozygous and have MAF between *f*_1_ and *f*_2_. Applying the same reasoning as above, we model the number of alleles with frequency between *f*_1_ and *f*_2_ inserted by gene conversion over *g* meioses and *l* base pairs as having a Poisson distribution with mean *lgθh*(0, *f*_2_)/2. The distribution is the same for alleles with frequency between 1 – *f*_2_ and 1 – *f*_1_.

We also consider genotype errors. Our method is concerned with low frequency alleles that are observed in two out of three IBD haplotypes. The probability of seeing this at any given base pair, conditional on the true alleles being identical in the three haplotypes, is defined to be for alleles with MAF ≤ *f*_1_ and be *ϵ*_2_ for alleles with MAF between *f*_1_ and *f*_2_. Thus for *l* base pairs, we model the numbers of such errors as having Poisson distributions with means *lϵ*_1_ and *lϵ*_2_, respectively.

The parameters *μ, θ, ϵ*_1_ and *ϵ*_2_ are not known a priori. We compute a likelihood across a grid of values for these parameters, and report the parameter values that maximize the likelihood. Our primary interest is in estimating *μ*.

### Modelling of IBD

We compute likelihoods for sets of three mutually-IBD haplotypes. We need to model the genealogical relationship between these three haplotypes.

In computing the likelihood, we will sum over the different possible relationships. Here we consider a single, generalizable relationship, 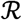, in which haplotypes A and B have their most recent common ancestor first, *g*_1_ generations ago, and the ancestor of A and B has its most recent common ancestor with C occurring *g*_1_ + *g*_2_ generations ago (Figure 1). The probability of this particular relationship depends on the population’s demographic history. If *N*[*g*] is the effective size of the population *g* generations before the present, then the probability of the relationship is^10^

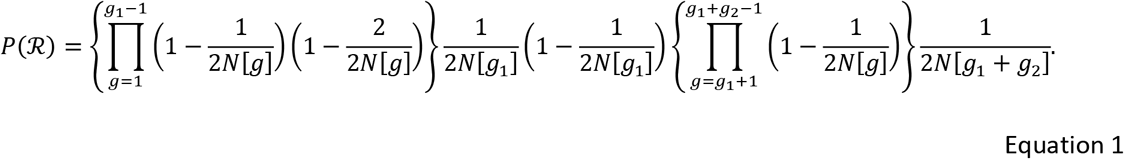

**Figure 1.**
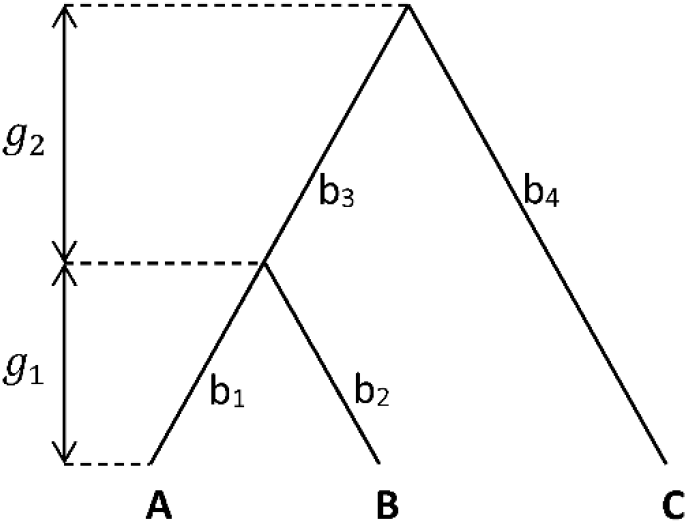
The genealogical relationship between three IBD haplotypes. A, B, and C are mutually-IBD haplotypes. Haplotypes A and B have a common ancestor *g*_1_ generations before the present, while C and the common ancestor of A and B have a common ancestor *g*_1_ + *g*_2_ generations before the present. The branches of the genealogical tree are labelled b_1_, b_2_, b_3_, and b_4_.

We use the distribution of lengths of pairwise IBD sharing in the sample to estimate the effective population sizes *N*[*g*] using the IBDNe program.^13^

The probability of the lengths of the observed IBD segments between the three haplotypes, given the relationship can be found in Table S1, with derivations in Tian et al.^10^

### Probability of allele counts given relationship

For alleles with frequency < *f*_1_, and for alleles with frequency between *f*_1_ and *f*_2_, we count the number of instances in which haplotypes A and B carry the allele but C does not, the number of instances in which haplotypes A and C carry the allele but B does not, and the number of instances in which haplotypes B and C carry the allele but A does not. These counts apply to the region shared IBD by all three haplotypes after trimming 0.5 cM from each end. We write *l_I_* for the base pair length of this region. The trimming accounts for uncertainty in the exact endpoints of the IBD segment.

Given the relationship in Figure 1, with branches labelled as in the figure, the low-frequency variants carried by A and B but not C can be obtained through one of the following ways. 1) A mutation on branch b3. 2) A gene conversion on branch b3 that replaces the high-frequency allele with the low-frequency allele. 3) A gene conversion on branch b4 that replaces the low-frequency allele with the high-frequency allele. 4) Genotype error. Other possibilities involve low probability events such as two or more gene conversions at the same site, or back-mutation, and we ignore these.

Given detectable IBD (e.g. length > 2 cM) between the haplotypes, their time to most recent common ancestor (*g*_1_ + *g*_2_) is within the past few hundred generations, and the frequency of any allele created by mutation on branch b_3_ will be low in the population. Under our model, the number of alleles with frequency < *f*_1_ that are shared by A and B but not C follows a Poisson distribution with mean

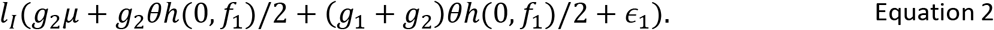

Under our model, the number of alleles with frequency between *f*_1_ and *f*_2_ that are shared by A and B but not C follows a Poisson distribution with mean

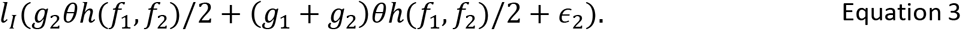

Given the relationship in Figure 1, with branches labelled as in the Figure, the low-frequency variants carried by A and C but not B can be obtained through one of the following ways. 1) A gene conversion on branch b2 that replaces the high-frequency allele with the low-frequency allele. 2) Genotype error. Other possibilities involve low probability events such as two or more gene conversions at the same site, or back-mutation, and we ignore these. Thus the number of alleles with frequency < *f*_1_ that are shared by A and C but not B follows a Poisson distribution with mean

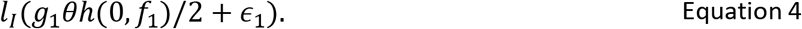

Similarly, the number of alleles with frequency between *f*_1_ and *f*_2_ that are shared by A and C but not B follows a Poisson distribution with mean

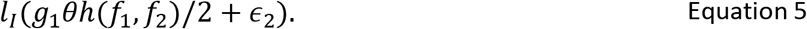

These same distributions apply to counts of alleles shared by B and C but not A.

### The likelihood

The observed data for each instance of three-way IBD sharing are the allele counts for the three haplotypes within the region shared IBD by all three haplotypes after trimming, the trimmed three-way IBD segment length measured in base pairs, and the untrimmed lengths of the pairwise IBD segments measured in cM. The likelihood (of *μ, θ, ϵ*_1_, *ϵ*_2_) is obtained by summing over the possible genealogical relationships, and for each one multiplying the prior probability of the relationship (Equation 1), the probability of the IBD lengths given the relationship (Table S1), and the probability of the allele counts given the relationship (Equations 2–5).

This gives the likelihood based on data from one instance of three-way IBD sharing. We find all instances of three-way IBD sharing, and multiply the likelihoods (or add the log-likelihoods) to obtain the overall likelihood. This is a composite likelihood because the instances of three-way IBD sharing are not fully independent. We perform a grid search to find the values of the parameters that maximize the likelihood. Confidence intervals are found by bootstrap resampling of the instances of three-way IBD sharing. We obtain 10,000 bootstrap estimates and report the 2.5^th^ and 97.5^th^ percentiles as the 95% confidence interval.

In order to reduce the computation time for the grid search, we use a three-stage approach. In the first stage, we use alleles with MAF < *f*_1_ and Equations 2 and 4 to obtain estimates of *μ, θ* and *ϵ*_1_. We discard the estimates for *μ, θ* from this analysis because much of the information about *θ* is contained in the higher frequency data, and because *μ* and *θ* are somewhat confounded without the use of the higher frequency data. We retain the estimate 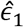 of *ϵ*_1_, for use in the third stage. In the second stage, we use alleles with MAF between *f*_1_ and *f*_2_ and Equations 3 and 5 to obtain estimates of *θ* and *ϵ*_2_. As in the first stage, we discard the estimate for *θ* since it is based on partial data, but we retain the estimate 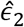 of *ϵ*_2_. In the third stage, we fix *ϵ*_1_ and *ϵ*_2_ at 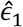 and 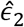, and use alleles with MAF < *f*_1_ as well as alleles with MAF between *f*_1_ and *f*_2_ and Equations 2–5 to obtain overall estimates of the mutation rate *μ* and gene conversion rate *θ*. For each parameter, we start the grid search with a wide range and large step size. If the resulting confidence interval falls within the range, we then reduce the range and the step size. This iterative step is repeated until the desired level of precision is achieved. We provide further details of the procedure for performing the three-stage likelihood maximization in the context of using the supplied java program in Supplemental Methods.

### Accounting for phase uncertainty

Common variants can be phased extremely accurately with statistical methods in large samples.^14^ We use the statistically-inferred phase for variants with minor allele frequency (MAF) ≥ 1%. Because lower frequency variants can’t be phased as well using standard statistical phasing, we use a different approach for variants with MAF < 1%.

Here we are considering alleles that are rare (frequency < 1%) yet are carried by two individuals who share a haplotype IBD. The question is whether the rare allele is on the IBD haplotype or not. In most cases it will be on the IBD haplotype because if it is not then the probability of seeing this rare allele in both individuals is small. Yet if there is evidence for one or both individuals that the allele may be coming in through the individual’s other (non-IBD) haplotype, we exclude it from the count.

Identification of IBD segments is performed using the statistically phased haplotypes, and segments are identified with respect to the haplotype indices. For example, we may find that at one position, individuals A and B are identical by descent on A’s haplotype 1 and B’s haplotype 2. We may also observe that A and D are identical by descent at the same position, but with A’s haplotype 2 and D’s haplotype 1. Thus, in this case A’s IBD sharing with D is not on the same haplotype as A’s IBD sharing with B.

Consider individuals A and B who have a haplotype that is also shared with C (three-way IBD sharing). For the sake of concreteness, suppose it is haplotype 1 of A and haplotype 1 of B that is IBD with haplotype 1 of C. Now suppose we observe that A and B both carry one copy of a certain rare allele “z” at a position that is within the region of three-way sharing, but that C does not carry this allele. We want to know whether the allele is on haplotype 1 of A and haplotype 1 of B and hence should be counted in the likelihood calculations. Since this allele is rare, we do not have a direct estimate of its phase. We look to see if there is any individual D who has one or more copies of “z”, who does not have a haplotype identical by descent with haplotype 1 of A or with haplotype 1 of B, and who has a haplotype identical by descent with haplotype 2 of A or with haplotype 2 of B at the position. We require that such IBD must extend at least 0.5 cM on either side of the position to protect against possible mis-estimation of the IBD segment endpoints. If we find one or more such individuals, we determine that the rare variant is on haplotype 2 of A and B, and is thus not part of the three-way IBD with C and can be ignored in the likelihood calculation. A special case that will occasionally occur is when A and/or B is homozygous for “z”. In this case, we always include the variant in the likelihood calculation.

### Simulated data

We used MaCS^15^ to simulate a dataset with a gene-conversion initiation rate of 2.0 × 10^−8^ per base pair per meiosis and mean gene conversion tract length of 300bp, which is close to previously reported estimates of gene conversion rate using human data.^16^ The mutation rate for the simulation was 1.3 × 10^−8^ per base pair per meiosis, and the recombination rate was 10^−8^ per base pair per meiosis. The simulation sample size was 2000 diploid individuals, and 30 chromosomes of length 100 Mb were simulated. The simulated demographic history was the ‘European-American model’ described in Tian et al.,^10^ which is based on an IBDNe^13^ analysis of TOPMed Framingham Heart Study data.^17^ We added genotype error to each simulated single nucleotide variant (SNV), changing each allele with probability 0.01%.

We inferred haplotype phase with Beagle 5.1^18^ for variants with MAF > 1%. We inferred IBD segments using hap-IBD^19^ with SNVs with MAF ≥ 0.25 (=*f_2_*), the minimum seed length (min-seed) set to 0.5 cM, the minimum extension length (min-extend) set to 0.1 cM, and the maximum gap length (max-gap) set to 5000 base pairs. Three-way IBD was obtained using segments of length 2.5 – 6 cM. We inferred the recent effective population size with IBDNe^20^ using the inferred IBD segments and default settings.

### TOPMed data

We estimated mutation rates using TOPMed project whole genome sequence data^17^ from 4,166 European-descent individuals in the Framingham Heart Study (FHS, dbGaP: phs000974.v4.p3), 2,996 White non-Hispanic individuals in the My Life Our Future project (MLOF, dbGaP: phs001515.v2.p2), and 1,586 Black non-Hispanic individuals in the TOPMed Hypertension Genetic Epidemiology Network Study (HyperGEN, dbGaP: phs001293.v2.p1). For these data, we used haplotype phase inferred in a previous analysis with a larger set of TOPMed project individuals.^21^ We used the deCODE genetic map^22^ throughout the analysis.

We used hap-IBD^19^ to detect pairwise IBD segments from the phased haplotypes using the same MAF filtering and parameters as for the simulated data. We used ibd-ends^23^ for a second step of IBD segment inference because we previously found that in real sequence data large spikes in IBD rate occur at some locations,^10; 23^ and the application of ibd-ends solves this issue.^23^ For the ibd-ends analysis, we used the output from the hap-IBD analysis as the input IBD segments, and we kept all parameters at their default values. The ibd-ends program estimates the posterior distribution of the endpoints for each IBD segment and returns the posterior medians which we used as the adjusted endpoints of the IBD segments.

When searching three-way IBD sharing, we restricted to only IBD segments with length between 2.5 cM to 6 cM as in the simulated data. We also excluded any IBD segments from duplicated samples, identical twins, and parent-offspring pairs. We identified such pairs of individuals from the provided pedigree file if available, or otherwise from the degree of relatedness estimated by IBDkin.^24^

In the FHS cohort we removed from the analysis the IBD segments shared between 2,415 pairs of individuals who are identified as duplicated samples, monozygotic twins, or parent and offspring in the provided pedigree. Among White non-Hispanic individuals in the MLOF cohort, we removed the IBD segments shared between 385 pairs of 1^st^ degree relatives and 8 pairs of 0^th^ degree relatives identified by IBDkin^24^. In the HyperGEN Black non-Hispanic cohort, IBDkin^24^ found 803 pairs of 1^st^ degree relatives and 2 pairs of 0^th^ degree relatives, and we removed the IBD segments shared between these pairs. The identified three-way IBD in the FHS cohort covered 2.75 gigabases across the autosomes. In the White non-Hispanic participants in the MLOF study, the identified three-way IBD sharing covered 2.73 gigabases across the autosomes. In the HyperGEN Black non-Hispanic cohort, the identified three-way IBD covered 2.55 gigabases across the autosomes. The reduced IBD coverage for the HyperGEN cohort is primarily due to the smaller sample size.

We inferred the recent effective populations sizes of the three study populations using the default settings of IBDNe^20^ with the IBD segments output by ibd-ends. We estimated likelihoods for each study across a search grid as for the simulated data, and multiplied the likelihoods to obtain overall likelihoods. We report the mutation rate that maximizes the likelihood separately for each study, and overall for the combined analysis.

## Results

### Analysis of simulated data

In the simulated data, which has a mutation rate of 1.3 × 10^−8^ per base pair per meiosis, we obtained a mutation rate estimate of 1.31 × 10^−8^ per base pair per meiosis with a 95% confidence interval of [1.28 × 10^−8^, 1.33 × 10^−8^]. If using the true (simulated) phase rather than estimated phase, the estimate is similar at 1.31 × 10^−8^ per base pair per meiosis with a 95% confidence interval of [1.29 × 10^−8^,1.34 × 10^−8^]. In contrast, when using estimated phase but ignoring the phasing uncertainty, the estimated mutation rate is 1.21 × 10^−8^ per base pair per meiosis with a 95% confidence interval of [1.19 × 10^−8^, 1.23 × 10^−8^]. Hence the simulation results demonstrate that the proposed method can effectively adjust for the phasing uncertainty.

We also compared the estimated gene conversion rates to the simulated gene conversion rate, which is 2.0 × 10^−8^ × 300 = 6.0 × 10^−6^. With true haplotype phase, the estimated gene conversion rate is 6.3 × 10^−6^ with a 95% confidence interval of [6.1 × 10^−6^, 6.4 × 10^−6^]. With inferred haplotype phase, the estimated gene conversion rate is 4.8 × 10^−6^ with 95% confidence interval of [4.6 × 10^−6^,4.9 × 10^−6^]. This downward bias cannot be corrected by increasing the allele frequency threshold for IBD detection, and is likely due to the effect of gene conversion events on phasing accuracy. Thus, in real population data, for which true phase is not available, the gene conversion rate cannot be accurately estimated. Nevertheless, this does not appear to affect the accuracy of the estimation of mutation rate.

### Analysis of TOPMed data

The joint estimate of mutation rate using data from the three TOPMed cohorts is 1.24 × 10^−8^ per base pair per generation with 95% confidence interval [1.18 × 10^−8^,1.33 × 10^−8^].

Using the FHS cohort alone, the estimated mutation rate is 1.28 × 10^−8^ per base pair per meiosis with 95% confidence interval [1.21 × 10^−8^,1.38 × 10^−8^]. Using the White non-Hispanic MLOF cohort we obtain a mutation rate estimate of 1.14 × 10^−8^ per base pair per meiosis with 95% confidence interval [1.04 × 10^−8^, 1.27 × 10^−8^]. Using the Black non-Hispanic HyperGEN cohort we estimate the mutation rate to be 1.34 × 10^−8^ per base pair per generation with 95% confidence interval [1.22 × 10^−8^,1.47 × 10^−8^]. The confidence intervals from all three studies overlap (Figure 2), indicating that the results are consistent with all three populations having the same mutation rate.

**Figure 2.**
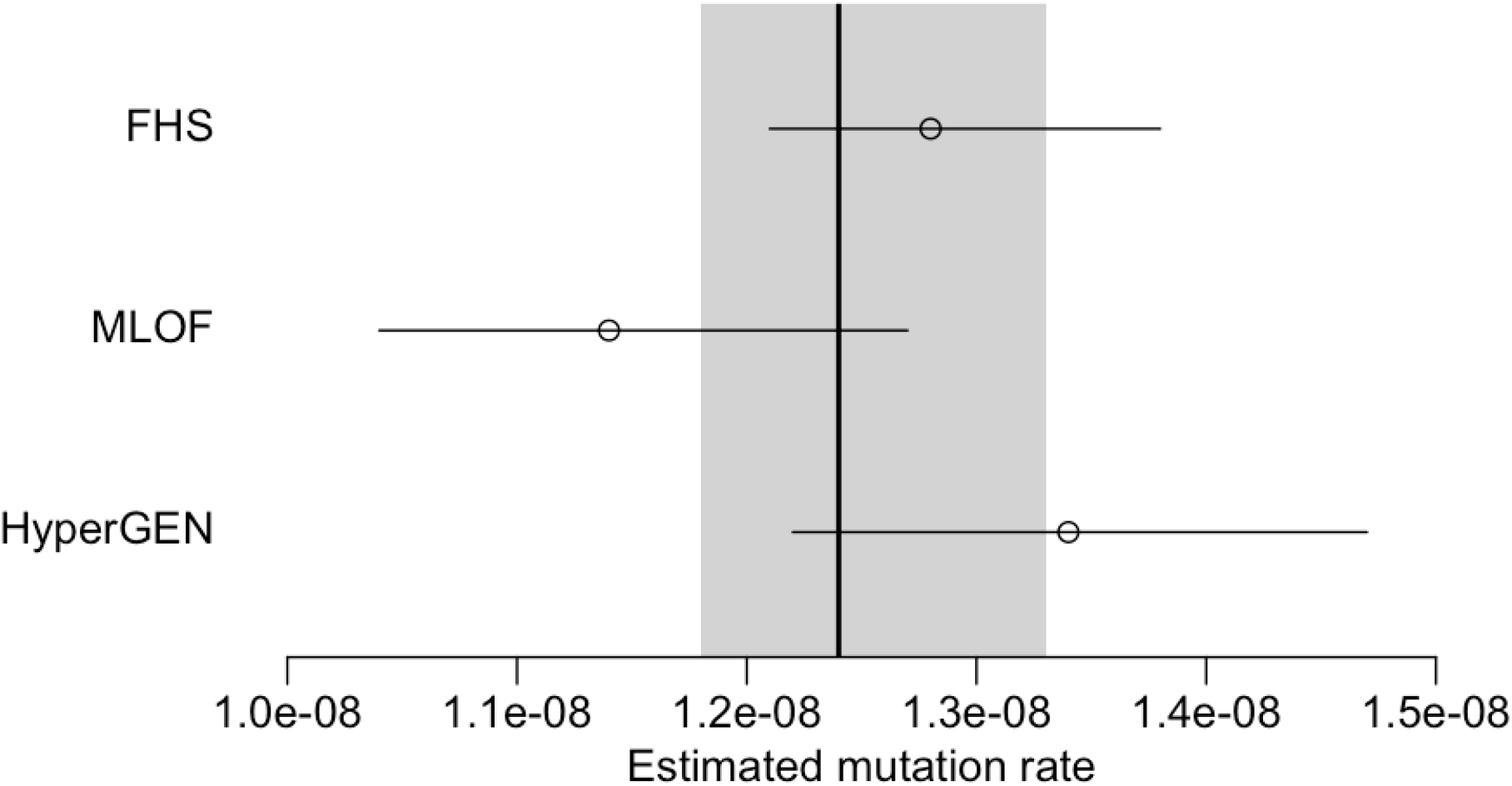
Estimated mutation rates in the TOPMed FHS, MLOF, and HyperGEN cohorts. Point estimates for each dataset are shown along with 95% confidence intervals (horizontal lines). The overall estimate from joint analysis of all three datasets is shown by the vertical line, with the 95% confidence interval represented by the shaded area.

## Discussion

Our work in this paper utilizes the three-way IBD framework of Tian et al.^10^ This three-way IBD framework allows for very strong control over genotype error, by requiring that potential mutations be observed in two of three IBD individuals. The approach of Tian et al. required that haplotypes be accurately phased, including phasing of rare variants, which in practice requires that the sequenced individuals include close relatives to enable the application of Mendelian phasing. Population samples can be statistically phased using linkage disequilibrium-based methods such as Beagle,^21^ however very rare variants tend not to be phased well with such methods. We thus developed a method for counting mutations on the IBD haplotypes when the underlying individual data is from population samples with uncertain phase. Again the three-way framework is very helpful in this mutation counting process because it largely removes the effect of allele errors on the non-IBD haplotypes of the individuals, because it removes the most recent mutations on the non-IBD haplotypes (which are much less likely to be shared by two of the three individuals), and because the additional “D” haplotypes that we introduce for the phasing can be IBD with the non-IBD haplotype of either of two individuals which increases the chance of finding such a haplotype.

Additionally, we incorporated the effects of gene conversion directly into the likelihood rather than accounting for them with a post-processing regression on maximum allele frequency.^8; 10^ This has the potential to increase statistical efficiency in estimating the mutation and gene conversion rates. However, we found that the method underestimates the gene conversion rate in analysis of population data. This is likely because gene conversion can induce phasing errors in common variants that disrupt the detection of IBD segments from common variants. Thus, a proportion of IBD segments involving gene conversion are lost to analysis. In contrast, mutation rates are still accurately estimated. Mutation rates are based on rare variants that are not included in the IBD detection process.

We analyzed data from more than 8000 individuals from three TOPMed studies including European-American and African-American populations and found that estimated mutation rates were not statistically different between the analyzed populations. The overall estimated genome-wide average mutation rate for SNVs was 1.24 × 10^−8^ per base pair per generation. This rate is consistent with our previous IBD-based estimates, but has tighter confidence intervals due to the larger sample size enabled by the new methodology presented here.^10^ Our estimated rate is higher than some estimates based on direct ascertainment of de novo variants in parent-offspring trio data,^3; 25^ with possible reasons including the stringent filtering that is needed in trio analyses in order to avoid counting false-positive mutations.^3; 4; 25^ A caveat of our results is that they apply only to SNVs, and only to those parts of the autosomes that are well-covered by called genotypes, which is approximately 2.7 gigabases in the data that we analyzed.

## Supplemental Data

Supplemental data consist of 1 table and Supplemental Methods.

## Acknowledgments

The methodological and analytical work performed in this study was supported by R01 HG005701 from the National Human Genome Research Institute (NHGRI). The content is solely the responsibility of the authors and does not necessarily represent the official views of the National Institutes of Health. Molecular data for the Trans-Omics in Precision Medicine (TOPMed) program was supported by the National Heart, Lung and Blood Institute (NHLBI). Core support including centralized genomic read mapping and genotype calling, along with variant quality metrics and filtering were provided by the TOPMed Informatics Research Center (3R01HL-117626-02S1; contract HHSN268201800002I). Core support including phenotype harmonization, data management, sample-identity QC, and general program coordination were provided by the TOPMed Data Coordinating Center (R01HL-120393; U01HL-120393; contract HHSN268201800001I). We gratefully acknowledge the studies and participants who provided biological samples and data for TOPMed. The Framingham Heart Study (FHS) was supported by contracts NO1-HC-25195 and HHSN268201500001I from the National Heart, Lung and Blood Institute (NHLBI) and grant supplement R01 HL092577-06S1; genome sequencing was funded by HHSN268201600034I and U54HG003067. The My Life, Our Future samples and data are made possible through the partnership of Bloodworks Northwest, the American Thrombosis and Hemostasis Network, the National Hemophilia Foundation, and Bioverativ; genome sequencing was funded by HHSN268201600033I and HHSN268201500016C. The Hypertension Genetic Epidemiology Network Study is part of the NHLBI Family Blood Pressure Program; collection of the data represented here was supported by grants U01 HL054472, U01 HL054473, U01 HL054495, and U01 HL054509; genome sequencing was funded by R01HL055673.

## Web Resources

Software for mutation rate estimation: https://github.com/tianxiaowen/mutation_unphased

Beagle: https://faculty.washington.edu/browning/beagle/beagle.html

hap-IBD: https://github.com/browning-lab/hap-ibd

ibd-ends: https://github.com/browning-lab/ibd-ends

MaCS: https://github.com/gchen98/macs

---

**Table S1:**
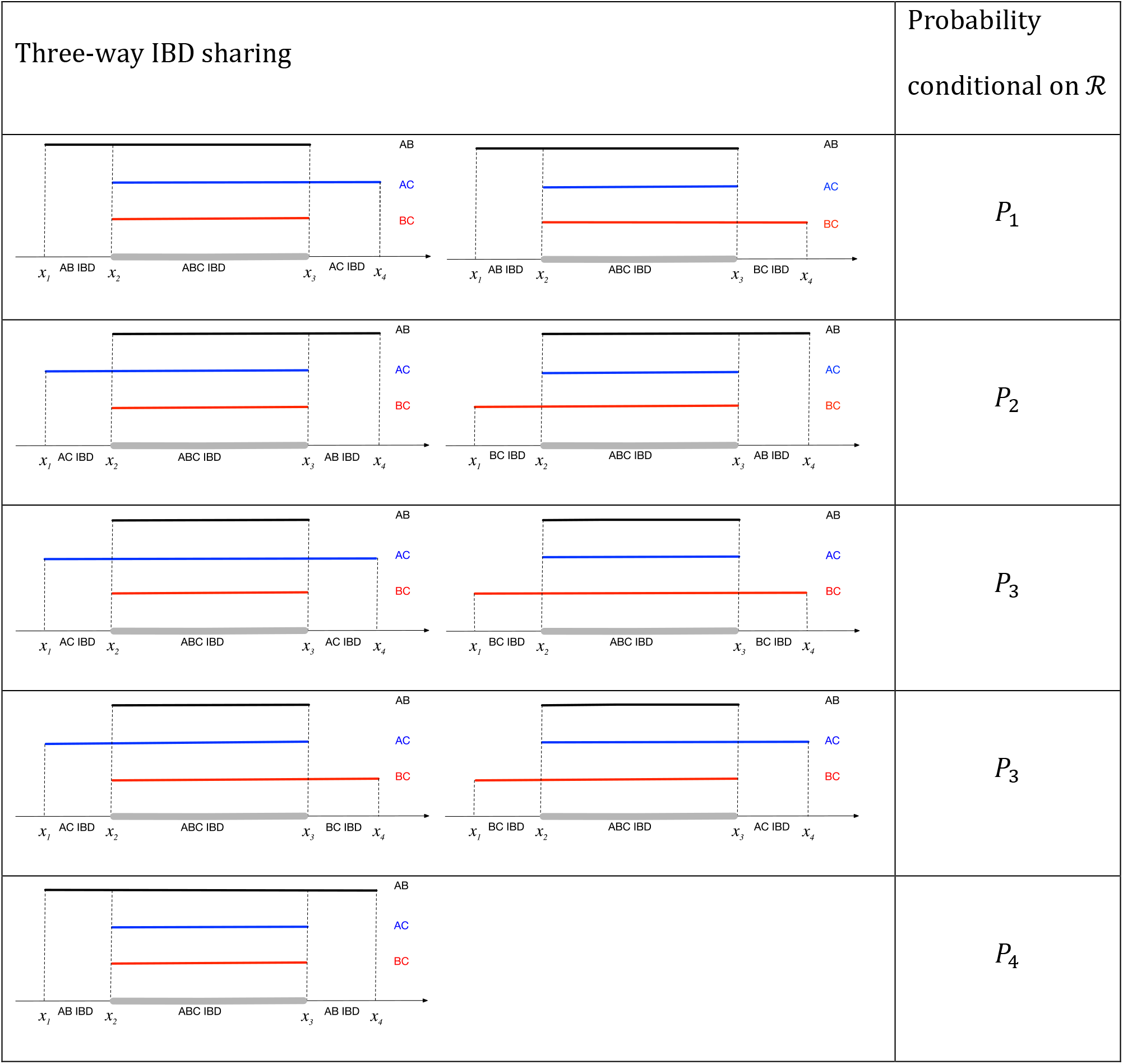
Probabilities for all possible three-way IBD segment configurations. (Adapted from Table S1 in Tian et al. 2019.) The right column gives the probability of the IBD segment configuration in the left column, conditional on the relationship 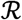 shown in Figure 1 (haplotypes A and B coalesce generations ago and their common ancestor and haplotype C coalesce *g*_1_ + *g*_2_ generations ago). The corresponding probability is given by one of the equations represented by *P*_1_, *P*_2_, *P*_3_, *P*_4_ below. The positions *x*_1_, *x*_2_, *x*_3_, *x*_4_ of changes in IBD status are measured in Morgans.

Let *G*_1_(*x*; *λ*) = *λe*^−*λx*^ denote an exponential distribution and 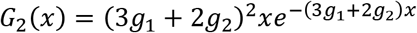 denote the gamma distributions with shape parameter 2 and rate parameter 3*g*_1_ + 2*g*_2_. Then

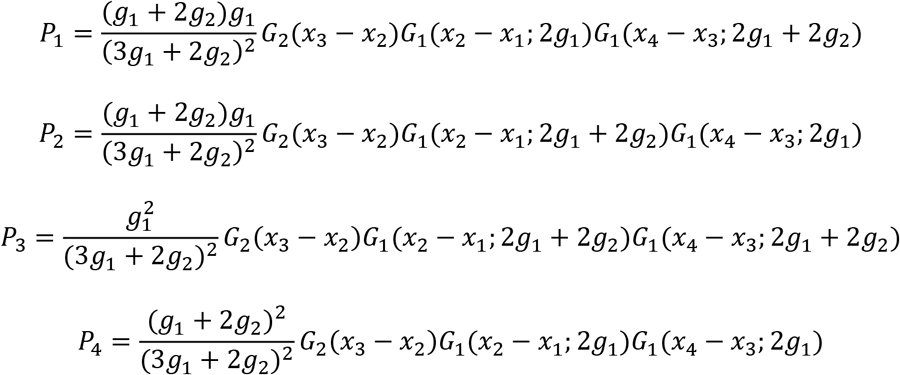

## Supplemental Methods

### Two-stage iterative grid search for maximum likelihood estimates with the program addGC.v3.jar

#### Stage 1: use alleles with MAF < *f*_1_ and Equations 2 and 4 to obtain estimates of *μ, θ* and *ϵ*_1_

1.1. Set stage = 1 and start the grid search with wide ranges and large step sizes for *μ, θ* and *ϵ*_1_. For example:
  *μ*: mu.start=1E-8 mu.end=2E-8 mu.step=5E-10
  *θ*: gc.start=1E-6 gc.end=5E-6 gc.step=1E-6
  *ϵ*_1_: err1.start=1E-15 err1.end=1E-6 err1.ratio=5 When the stage is set to 1 in addGC.v3.jar, *ϵ*_2_ will be set to 0 automatically.
1.2 If the resulting bootstrap confidence interval falls within the range for each parameter, then the range and the step size can be reduced. Otherwise, increase the range of search. In the analysis of simulated datasets, the final search step size for *ϵ*_1_ was set to err1.ratio=1.5.
1.3 Retain the estimate 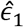 of *ϵ*_1_.

#### Stage 2: use alleles with MAF between *f*_1_ and *f*_1_ and Equations 3 and 5 to obtain estimates of *θ* and *ϵ*_2_

2.1. Set stage = 2 and start the grid search with wide ranges and large step sizes for *θ* and *ϵ*_1_. For example:

*θ*: gc.start=1E-6 gc.end=5E-6 gc.step=1E-6
*ϵ*_2_: err2.start=1E-15 err2.end=1E-6 err2.ratio=5 When the stage is set to 2 in addGC.v3.jar, *μ* and *ϵ*_1_ will be set to 0 automatically.
2.2 If the resulting bootstrap confidence interval falls within the range for each parameter, then the range and the step size can be reduced. Otherwise, increase the range of search. In the analysis of simulated datasets, the final search step size for *ϵ*_2_ was set to err2.ratio=1.5.
2.3 Retain the estimate 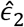 of *ϵ*_2_.

#### Stage 3: fix *ϵ*_1_ and *ϵ*_2_ at 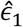 and 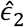, and use alleles with MAF < *f*_1_ as well as alleles with MAF between *f*_1_ and *f*_2_ and Equations 2–5 to obtain overall estimates of the mutation rate *μ* and gene conversion rate *θ*.

3.1. Set stage = 3 and start the grid search with wide ranges and large step sizes for *μ* and *θ* while setting the same starting and ending points for *ϵ*_1_ and *ϵ*_2_ at 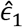 and 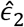, respectively. For example:
  *μ*: mu.start=1E-8 mu.end=2E-8 mu.step=5E-10
  *θ*: gc.start=1E-6 gc.end=5E-6 gc.step=1E-6
  *ϵ*_1_: err1.start=4.379E-9 err1.end=4.379E-9
  *ϵ*_2_: err2.start=3.464E-14 err2.end=3.464E-14
3.2 If the resulting bootstrap confidence interval falls within the range for each parameter, then the range and the step size can be reduced. Otherwise, increase the range of search. In the analysis of simulated datasets, the final search step size for *μ* and *θ* were set to mu.step=1E-10 and gc.step=0.05E-6, respectively.
3.3 Retain the final estimates of 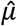 and 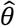.

